# An *Erwinia amylovora* inducible promoter for improvement of apple fire blight resistance

**DOI:** 10.1101/767772

**Authors:** Gaucher Matthieu, Righetti Laura, Aubourg Sébastien, Dugé de Bernonville Thomas, Brisset Marie-Noёlle, Chevreau Elisabeth, Vergne Emilie

## Abstract

Intragenesis is an important alternative to transgenesis to produce modified plants containing native DNA only. A key point to develop such a strategy is the availability of regulatory sequences controlling the expression of the gene of interest. With the aim of finding apple gene promoters either inducible by the fire blight pathogen *Erwinia amylovora (Ea)* or moderately constitutive, we focused on polyphenoloxidase genes (*PPO*). These genes encode oxidative enzymes involved in many physiological processes and have been previously shown to be up-regulated during the *Ea* infection process. We found ten PPO and two PPO-like sequences in the apple genome and characterized the promoters of *MdPPO16* (*pPPO16*) and *MdKFDV02 PPO*-like (*pKFDV02*) for their potential as *Ea*-inducible and low-constitutive regulatory sequences respectively. Expression levels of reporter genes fused to these promoters and transiently or stably expressed in apple were quantified after various treatments. Unlike *pKFDV02* which displayed a variable activity, *pPPO16* allowed a fast and strong expression of transgenes in apple following *Ea* infection in a Type 3 Secretion System dependent manner. Altogether our results indicate that *pKFDV02* did not keep its promises as a constitutive and weak promoter whereas *pPPO16*, the first *Ea*-inducible promoter cloned from apple, can be a useful component of intragenic strategies to create fire blight resistant apple genotypes.

**Key message:** *pPPO16*, the first *Ea*-inducible promoter cloned from apple, can be a useful component of intragenic strategies to create fire blight resistant apple genotypes.

## Introduction

*Erwinia amylovora* (*Ea*) is a necrogenic enterobacterium causing progressive necrosis on flowers and succulent shoots in members of the *Malinae* tribe of the *Rosaceae* family including apple (*Malus x domestica* Borkh; Vanneste 2000). Rapid invasion of the bacteria into branches and trunks can lead to the death of the trees within a growing season for the most susceptible cultivars. At the cellular level, the bacteria use a Type 3 Secretion System (T3SS) to deliver effectors into the plant cells, to induce membrane disruption and oxidative burst leading to cell death (Vrancken et al. 2013). H_2_O_2_ is one of the first detectable ROS (Reactive Oxygen Species) produced during this infection process (Vrancken et al. 2013). Fire blight outbreaks are sporadic, particularly difficult to control and improving host resistance is by far the most effective option to control the disease (Paulin, 1996). Breeding for Fire blight resistance is therefore an active area of research with the identification of genetic resistance factors including quantitative traits loci (Khan et al. 2012), a “resistance” gene (R gene) implicated in pathogen recognition (*FB_MR5*; Vogt et al., 2013) or defense mechanisms downstream recognition (Vrancken et al. 2013).

Numerous attempts to create fire blight resistant apple transgenic lines have been performed with various degrees of success. For example, a number of studies were based on the expression of foreign genes encoding insect lytic proteins (Borejsza-Wysocka et al. 2010), a chalcone 3-hydroxylase (Hutabarat et al. 2016), a viral EPS-depolymerase (Flachowsky et al. 2008a) or the *Ea* HrpN protein (Vergne et al. 2014). Other studies tested the introgression of the fire blight resistance gene *FB_MR5* (Broggini et al. 2014; Kost et al. 2015), the overexpression of apple defense genes such as *MpNPR1* (Malnoy et al. 2007), *MbR4* R gene (Flachowsky et al. 2008b), or the silencing or gene editing of potential apple susceptibility factors such as *HIPM* (Malnoy et al. 2008; Campa et al. 2019), *DIPM* (Pompili et al. 2020) or *FHT* (Flachowsky et al. 2012). To our knowledge, the only cisgenic strategy employed to improve fire blight resistance of apple was performed with the *FB_MR5* resistance gene controlled by its native regulatory sequences (Kost et al. 2015).

Intragenesis and cisgenesis are alternatives to transgenesis defined by Rommens et al. (2007) and Schouten et al. (2006) respectively, and are based on the exclusive use of genetic sequences from the same (or a sexually compatible) species. These strategies aim at improving crop breeding while considering the public’s reluctance toward the use of foreign genes usually present in the genetically modified plant varieties. In the case of cisgenesis, coding sequences (CDS) must be in a sense orientation and flanked by their native promoter and terminator sequences, while intragenesis allows a reorganization of both regulatory and coding sequences, as well as the introduction of mutations (e.g., nucleotide substitutions, sequence deletions, duplications and inversions), to fine tune the expression of the CDS of interest (Holme et al. 2013). These techniques are of particular interest for perennial vegetatively propagated crops such as apple for which conventional breeding is very time-consuming (Limera et al. 2017). In addition, the selectable marker gene is eliminated from cisgenic as well as from intragenic plants, thus allowing sequential introduction of a new transgene, using the same selectable marker, in an elite variety (Halpin 2005).

The generation of intragenic/cisgenic apple plants requires the development and combination of different strategies. The selection of transgenic lines can be based on alternative selectable marker genes from apple such as genes implicated in anthocyanin production (Kortstee et al. 2011) or genes of which certain mutation gives resistance to herbicide (acetolactate synthase; Yao et al. 2013). A recombinase-mediated removal of the unwanted selectable marker sequence has also been used (Herzog et al. 2012; Righetti et al. 2014; Kost et al. 2015).

As for the regulatory sequences, so far, only the apple Rubisco promoter has been used to obtain the constitutive and high expression of an intragene, the R gene *Rvi6*, to control apple scab caused by the fungi *Venturia inaequalis* (*Vi*, Joshi et al. 2011). Overexpression of genes downstream R genes in the defense pathways (i.e. regulators and defense genes) can lead to enhanced resistance but with an important energetic cost that might impede primary plant functions or create developmental disorders. For example, constant overexpression of master-switch genes like *NPR1* (Pieterse and Van Loon, 2004) can lead to lesion mimic phenotypes (Fitzgerald et al. 2004) and be detrimental to plant development (Gurr and Rushton, 2005). Overexpression of phytoalexins or other antimicrobial compounds at high level can also damage tissue integrity (Großkinsky et al. 2012). Therefore, the use of pathogen-inducible promoters to drive regulators of defense pathways, *PR* genes or toxic antimicrobial genes is a necessity (Gurr and Rushton, 2005). In order to create apple intragenic lines resistant to *Ea*, we were interested in two kinds of regulatory sequences: (i) an inducible promoter with a fast and strong induction after *Ea* infection and able to trigger the production of defense mechanisms in the right place at the right time against the bacteria and (ii) a constitutive promoter with a moderate expression level. Such a promoter could ensure the permanent presence of immune receptors such as pattern recognition ones or ones encoded by R genes, with minimal negative tradeoff effects. Previous results led us to investigate the family of polyphenol oxidases (PPO) for this purpose. This complex family of enzymes catalyzes the hydroxylation of monophenols and/or the oxidation of di-phenolic compounds into quinones (Pourcel et al. 2007). A high increase of global enzyme activity has been reported in apple after *Ea* infection (Skłodowska et al. 2011; Gaucher et al. 2013) and preliminary studies on gene expression by RT-qPCR revealed a clear differential induction of *PPO* genes - or set of genes - after infection (Dugé de Bernonville, 2009).

Here, we took advantage of the recent high-quality apple genome (Daccord et al. 2017) to fully describe the apple PPO family and to select individual genes with differential expression after *Ea* infection. After cloning, promoters of interest were fused to reporter genes and transiently or stably transformed in apple. This allowed the assessment of their activity under various stresses in order to evaluate their usefulness in future intragenesis strategies for apple resistance to *Ea*.

## Material and Methods

### Material, growth and inoculation conditions

#### Apple

Four *Malus x domestica* genotypes were used in this work: the ornamental cv. ‘Evereste’, the rootstock ‘MM106’ and the table apples ‘Golden Delicious’ and ‘Gala’. Experiments were performed in greenhouse on actively growing shoots of young grafts (‘Evereste’ and ‘MM106’) grafted on ‘MM106’, or on actively growing plants not grafted (‘Golden Delicious’), and grown under greenhouse conditions (natural photoperiod, temperatures between 17 and 22°C). Experiments were also performed on *in vitro*–growing shoots of three to four cm, used 4 weeks after rooting (Online Resource 1). Micropropagation conditions were as described in Righetti et al. (2014) and rooting conditions as previously reported (Faize et al. 2003).

### *Erwinia amylovora* culture, inoculation and experiments

Two *Ea* strains were used in this study: wild-type *Ea* CFBP1430 (*Ea* wt; Paulin and Samson, 1973) and PMV6023, a non-pathogenic T3SS-defective mutant of *Ea* wt, mutated in *hrcV* (*Ea* t3ss; Barny, 1995). Prior to each experiment, bacteria were subcultured at 26°C overnight on solid King’s medium B (King et al. 1954) supplemented with chloramphenicol (20 μg/mL) for the mutant. Bacterial inocula were prepared in sterile distilled water to yield a concentration of 10^7^ colony-forming units (CFU)/mL, supplemented with 0.01 % (v/v) of wetting agent Silwet (L-77, De Sangosse Ltd, Cambridge, UK). Mock corresponded to sterile water supplemented with the wetting agent Silwet.

For greenhouse growing plants inoculation was performed by vacuum infiltration as described in Pontais et al. (2008). Briefly, the top of growing shoots were submerged in bacterial suspension and the vacuum was applied for 2 min at −0.09 Mp (Online Resource 1).

In related experiments, leaf samples were immediately frozen in liquid nitrogen and kept at −80°C until analysis. Sampling concerned the youngest expanded leaf of each plant labeled the day of the inoculation. Each sample is a pool of leaves from three different plants and two (n=2; *PPO* genes expression analysis in ‘Evereste’ and ‘MM106’ genotypes) to three (n=3; promoters analysis in ‘Golden Delicious’ transgenic lines) biological repeats have been made by condition (genotype/transgenic line x treatment x time).

For *in vitro*–growing shoots, four weeks after rooting, shoots were separated from their roots, totally submerged in inoculum and vacuum infiltrated for 2 min at −0.09 Mp. Shoots were then dried on sterile filter paper and placed for 1 day back on micropropagation medium before sampling.

In related experiments, leaf samples were immediately frozen in liquid nitrogen and kept at −80°C until analysis. Sampling concerned all the leaves of each shoot. Each sample is a pool of leaves from three different plants and three (n=3; transient transformation assay on ‘Gala’ genotype) to six (n=6; *in vitro* experiments on ‘Golden Delicious’ transgenic lines) biological repeats have been made by condition (genotype/transgenic line x treatment x time).

### *Venturia inaequalis* culture, inoculation and experiment

The apple scab monoconidial isolate used was EU-B04 from the European collection of *V. inaequalis* from the European project Durable Apple Resistance in Europe (Lespinasse et al. 2000). Inoculum was prepared as described by Parisi and Lespinasse (1996) to obtain a final concentration of 2.5 × 10^5^ conidia/mL. Inoculation was performed as described by Parisi et al. (1993). Briefly, conidial suspension was applied to runoff on leaves with a manual sprayer. Plants were then incubated for two days under plastic tarpaulin and sprayed three times a day to assure constant leaf wetness. The tarpaulin was then removed and plants grew under greenhouse conditions. Mock corresponded to sterile water.

Leaf samples were immediately frozen in liquid nitrogen and kept at −80°C until analysis. Sampling concerned the youngest expanded leaf of each plant labeled the day of the inoculation. Each sample is a pool of leaves from three different plants and three biological repeats (n=3) have been made by condition (transgenic line x treatment x time).

### *Agrobacterium* culture

*Agrobacterium tumefaciens* EHA105 (Hood et al. 1993) containing binary expression vectors of interest (Online Resource 2) was cultured on LBA (LB Agar, Sigma-Aldrich, St. Louis, MO, USA) supplemented with appropriate antibiotics and incubated at 28°C for two days.

### H_2_O_2_ treatment

H_2_O_2_ (30% w/v solution, Fisher Scientific, Loughborough, UK) was used at 10 mM concentration on *in vitro*–growing shoots. Four weeks after rooting, shoots were separated from their roots and either cultured on micropropagation medium supplemented with 10 mM H_2_O_2_ during 1 day before sampling or vacuum infiltrated for 2 min at −0.09 Mp, dried on sterile filter paper and placed for 1 day back on micropropagation medium before sampling.

Leaf samples were immediately frozen in liquid nitrogen and kept at −80°C until analysis. Sampling concerned all the leaves of each shoot. Each sample is a pool of leaves from three different plants and six (n=6; *in vitro* experiments on ‘Golden Delicious’ transgenic lines) biological repeats have been made by condition (transgenic line x treatment x time).

### Transformation of apple

Agroinfiltration of *in vitro* rooted plants was used for transient transformation experiments. The inoculum for infiltration was a mix of the strain with the T-DNA of interest (Online Resource 2: *p35S, pKFDV02* or *pPPO16* from MM106) and the strain with the T-DNA carrying the gene coding the p19 protein of tomato bushy stunt virus as a suppresser of gene silencing (Voinnet et al. 2003), respectively at 5 ×10^8^ CFU/mL and 2.5 × 10^8^ CFU/mL. The cultures were re-suspended in induction buffer (10 mM MES pH 5.6, 10 mM MgCl2, 2% (w/v) sucrose and 150 μM acetosyringone) (Santos-Rosa et al. 2008), mixed at the desired concentration, incubated at 28°C with shaking for 3 h, and then supplemented with 0.002 % (v/v) of wetting agent Silwet before use. Four weeks after rooting, shoots were separated from their roots, totally submerged in inoculum and vacuum infiltrated for 2 min at −0.09 Mp. Shoots were then rinsed in 3 successive baths of sterile water, dried on sterile filter paper and placed for 6 days back on micropropagation medium without antibiotics before sampling.

Stable transformation experiments were carried out according as previously reported (Righetti et al. 2014). Presence of transgenes and absence of contaminating agrobacteria were monitored by PCR and sequencing of PCR products. Genomic DNA of apple leaves was extracted as described in Fulton et al. (1995). Primers used for the detection of (i) *A. tumefaciens* presence, (ii) *nptII* gene, (iii) *p35S:GUS* straddled amplification (iv) *pPPO16:GUS* straddled amplification, (v) *pKFDV02:GUS* straddled amplification and (vi) elongation factor 1α (*EF-1α*) coding gene as a marker of plant DNA suitability for PCR are available in Online Resource 3. Amplifications were performed using GoTaq^®^ Flexi DNA Polymerase (Promega, Madison, WI, USA) according to the manufacturer’s recommendations. The PCR reaction conditions were identical for the six genes except the hybridization step which was at 55°C and not 58°C for *A. tumefaciens* detection primers: 95°C for 5 min, followed by 35 cycles at 95°C for 30 s, 58°C for 45 s, 72°C for 1 min and 30 s, with a final extension at 72°C for 5 min. The PCR products were separated on a 2 % agarose gel. Transgenic lines and control plants were then propagated *in vitro* and acclimatized in a greenhouse as previously reported (Faize et al. 2003). Before acclimatization, the ploidy level of transgenic lines was checked by flow cytometry, as described in Chevreau et al. (2011), and tetraploid lines were eliminated.

### Characterization of apple PPO family

The annotated genes of the ‘Golden Delicious’ double haploid 13 genome (Daccord et al. 2017) have been screened for PFAM motifs specific to the PPO family, namely PF12142 and PF12143. The structural annotation of each detected locus was manually evaluated in considering BLASTX results and RNA contig alignments. The integrity of CDS has cautiously been checked in order to differentiate functional genes from pseudogenes. The twelve protein sequences deduced from complete and short CDS have been analyzed with targetP (Emanuelsson et al. 2007) and Predotar (Small et al. 2004) for the prediction of N-terminal targeting peptide for the plasts. Phylogenetic tree was built from full-length alignment with Neighbor-joining method, excluding gap positions and tested with Bootstrap method (Kumar et al. 2016). The percent identity matrix of CDS and proteins were built with MUSCLE (Edgar, 2004) and Clustal Omega (Sievers et al. 2011) respectively.

### Cloning of promoters

Sequence of CaMV 35S promoter in pK7WG2D (Karimi et al. 2002) and sequences of about 2 kb upstream *MdPPO16* (MD10G1299400) and *MdKFDV02* CDS (MD10G1298200) were downloaded. Primers for cloning (Online Resource 3) were designed with primer3plus (http://www.bioinformatics.nl/cgi-bin/primer3plus/primer3plus.cgi). Genomic DNA of apple ‘MM106’ was used as template for PCR amplification of promoters with a high fidelity DNA polymerase (Phusion Hot Start II DNA Polymerase, ThermoFisher Scientific, MA, USA) used according to the manufacturer instructions. Amplified fragments were then cloned into pGEM-T easy (Promega, Madison,WI, USA) or p-ENTR/D TOPO (Invitrogen, Carlsbad, CA, USA) when subsequent Gateway cloning was planned.

For apple stable transformation, promoters were cloned with the Gateway system via pENTR-D TOPO (Invitrogen, Carlsbad, CA, USA) into the destination vector pKGWFS7 (Karimi et al. 2002). In this vector the sequence under study controls the expression of a *GUS-GFP* reporter gene. 2219 bp upstream to *MdPPO16* and 2030 bp upstream to *MdKFDV02* start codons were cloned and the final constructs were transformed in *Agrobacterium* strain EHA105 with the helper plasmid pBBR-MCS5. As a positive control for stable transformation assays a plasmid pKGWFS7 carrying the CaMV 35S promoter was used (Online Resource 2).

For transient assays we used either the same plasmids as for stable transformation, or the binary vector pGREEN II 0800-LUC (Hellens et al. 2005). This vector is specifically designed to clone the sequence under study upstream to a firefly luciferase coding sequence. A renilla luciferase coding sequence under the control of a constitutive CaMV 35S promoter is also present as an internal control. The presence of the two luciferases in a single T-DNA reduces the intrinsic variability of leaf agroinfiltration and thus allows reproducible promoter activity quantification. Cloning into this vector was achieved by adding specific restriction sites to the primers. *Kpn*I and *Nco*I sites were added to the 5’ and 3’ ends respectively of *MdPPO16* and *MdKFDV02* promoters, while *Kpn*I and *Hind*III were added to primers used for CaMV 35S promoter. After digestion with restriction enzymes of both vectors and inserts, 1177 bp and 2030 bp of the sequences upstream the start codon were cloned for *MdPPO16* and *MdKFDV02*, respectively. As a positive control for transient assays 1027 bp of CaMV 35S promoter amplified from the plasmid pK7WG2D (Karimi et al. 2002) were also cloned. The final constructs were transformed in *Agrobacterium* strain EHA105 with the helper plasmid pSoup (Online Resource 2).

### DNA extraction, RNA extraction, reverse transcription, and gene expression analysis

Genomic DNA of leaves of apple ‘MM106’ was extracted as described in Fulton et al. (1995).

For RNA extraction, frozen leaves were ground to a fine powder in a ball mill (MM301, Retsch, Hann, Germany). RNA from leaves was extracted as described in Venisse et al. (2002). Purity and concentration of the samples were assayed with a Nanodrop spectrophotometer (ThermoScientific, Rockford, IL, USA). Reverse transcription was performed with M-MLV as described by Promega with OligodT 25 ng/μl or specific primers 0.04 μM final concentrations (Online Resource 3). Intron-spanning primers designed on the *EF-1α* gene were used to check the absence of genomic DNA contamination.

Quantitative PCR was used to quantify cDNA in samples. Briefly, 3.75 μL of the appropriately diluted samples (ranging from 4 to 12.5-fold) were mixed with 7.5 μL of qPCR mastermix (MasterMix Plus for SYBR© Green I with fluorescein, Eurogentec, Liège, Belgium) in a final volume of 15 μL. Primers designed with Primer3Plus were added according to their optimal concentration (determined for reaction efficiency near to 100%; calculated as the slope of a standard dilution curve; Pfaffl, 2001). Primer sequences are indicated in Online Resource 3. Reaction was performed on a DNA Engine thermal cycler Chromo4 (Bio-Rad, Hercules, CA, USA) using the following program: 95°C, 5 min; 35 cycles comprising 95°C 15 s, 60°C 45 s and 72°C 30 s with real-time fluorescence monitoring. Melt curves were performed at the end of each run to check the absence of primer-dimers and non-specific amplification products. Data were acquired and analyzed with MJ Opticon Monitor Software 3.1 (Bio-Rad, Hercules, CA, USA). Expression profiles of endogenous *PPO* genes were calculated using the 2^-ΔΔCt^ method and were corrected as recommended in Vandesompele et al. (2002), with three internal reference genes (*GADPH*, *TuA* and *Actin*) used for the calculation of a normalization factor. Data were transformed into log_2_ scale. Expression levels of the *GUS* and *FIRE* reporter genes were calculated using the 2^-ΔΔCt^ method and were corrected with the spectinomycin (*SPEC*) selection gene or the internal control *REN* respectively. *GUS* in pKGWFS7 did not possess an intron so in the transient assay this reporter gene actually dosed expression from both the plant and *Agrobacterium*. *SPEC* gene expression, specific from the bacteria because present in the plasmid but not in the T-DNA, was used to calibrate samples amongst themselves to eliminate the potential part of expression due to bacteria in the *GUS* measure differences.

### Luciferase and GUS activity assay

Frozen leaves were ground to a fine powder in a ball mill (MM301, Retsch, Hann, Germany). Luciferase activities were measured by using the dual luciferase assay system (Promega, Madison, USA) according to the manufacturer’s instructions but with some modifications. 150 μL of Passive Lysis Buffer were added to the resulting powders and samples were placed on ice for 15 min and vortexed several times in the meantime. For luciferase activity measurements (firefly and renilla), 10 μL of each extract were transferred into a 96-well white solid plate (Fisher Scientific ltd., Montreal, Quebec). The luminescence was measured using the FluoStar Optima Luminometer (BMG Lab Technologies, Offenburg, Germany) with the injection of 60 μL of LARII reagent (Firefly luciferase activity) and then 60 μL of the Stop & Glo reagent (Renilla luciferase activity). For Gus activity measurements, 800 μL of extraction buffer (50 mM Na_2_HPO_4_, pH 7.0, 1 mM Na_2_EDTA pH 8.0, 0.1% (v/v) Triton X-100, 0.1% (w/v) sodium lauryl sarcosine) were added to the leaf powders. The homogenates were centrifuged at 4°C for 1 min at 10000 rpm. The supernatants were 10-fold diluted and 100 μL of each dilution were transferred into a 96-well white solid plate. Quantitative fluorimetric GUS activity assay was performed using the FluoStar Optima Luminometer (BMG Lab Technologies, Offenburg, Germany) with the injection of 4 μL of the substrate 4-methylumbelliferyl β-glucuronide (MUG). Luciferase and GUS activities were standardized to the protein concentration (Bradford, 1976) of the extracts and firefly luciferase activity was normalized to renilla luciferase activity.

### Statistics analysis

All statistical analyses were performed with R 3.4 (R Development Core Team, 2016) by using the nonparametric rank-based statistical test Kruskal–Wallis. Treatments with significant influence (p<0.05) were studied more in depth by Fisher’s Least Significant Difference (LSD) as a *post hoc* test for pairwise comparisons (α = 0.05). Means with different letters are statistically significant.

## Results

### PPO encoding genes in the apple genome

Screening the ‘Golden Delicious’ double haploid 13 genome (Daccord et al. 2017) revealed the presence of a *PPO* gene family encompassing ten members with similar gene structure of one or two exons, encoding proteins ranging from 587 to 610 residues (Table 1). N-terminal signal peptides for chloroplast targeting were predicted for all of them. PPO proteins are characterized by three conserved PFAM domains: the tyrosinase superfamily domain PF00264, the PPOI_DWL domain PF12142 and the PPO1_KFDV domain PF12143. Apple *PPO* genes are located on two clusters on chromosomes 5 (five genes) and 10 (five genes), two chromosomes known to result from a whole genome duplication (Daccord et al. 2017). Close examination of these two chromosomal regions identified two additional PPO-like encoding genes, one on chromosome 5 and the other on chromosome 10, which conserved only the C-terminal KFDV domain and were also predicted to be addressed to the chloroplast (Table 1). Six pseudogenes were finally found, four on chromosome 5 and two on chromosome 10. Their CDS are disrupted by deletion, transposable element insertion, frameshift and/or stop codons (Table 1). We named *PPO* genes and pseudogenes *MdPPO01* to *MdPPO16*, and *PPO*-like genes *MdKFDV01* and *MdKFDV02*. Phylogeny based on 30 identified *PPO Rosaceae* homologs revealed five subfamilies (Fig. 1). Identity matrices obtained using nucleic or protein sequences of the ten apple *PPOs* showed a very high conservation level between accessions inside each apple *PPO* sub-family (Online Resource 4).

**Fig. 1.**
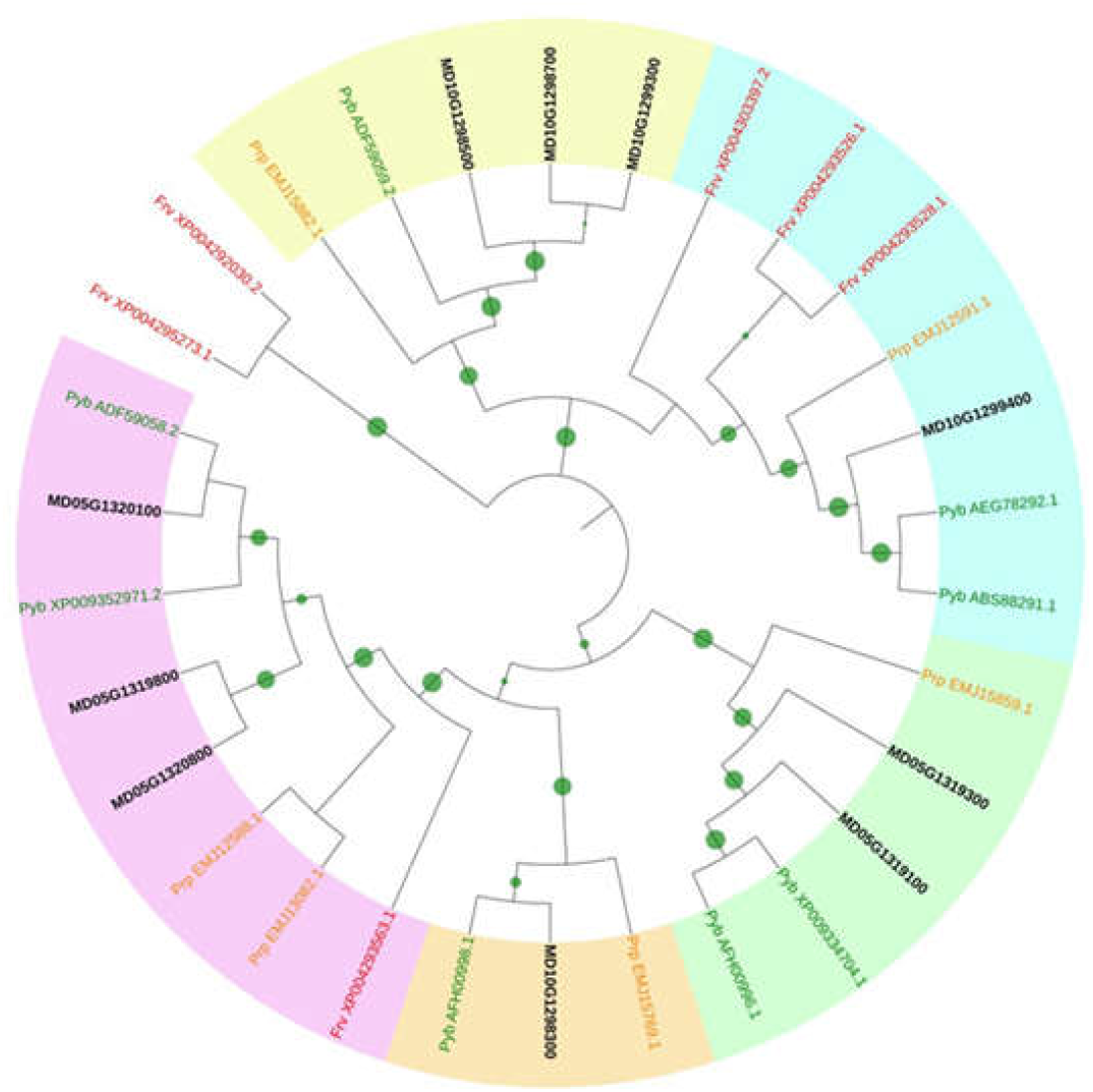
Phylogenetic tree of PPO homologs in *Rosaceae*. The tree was built with the neighbor-joining method from the multiple alignment of 30 homologous proteins. Gaps were ignored for tree building and 1000 bootstrap replicates were used to determine the robustness of each node (the bigger the green circle size, the more robust the node). Except for apple for which gene model ID is used, each protein is labeled with two letters (species) and its GenBank ID or XP number. Frv: Fragaria vesca, Prp: Prunus persica (L.) Batsch, Pyb: Pyrus bretschneideri

**Table 1.**
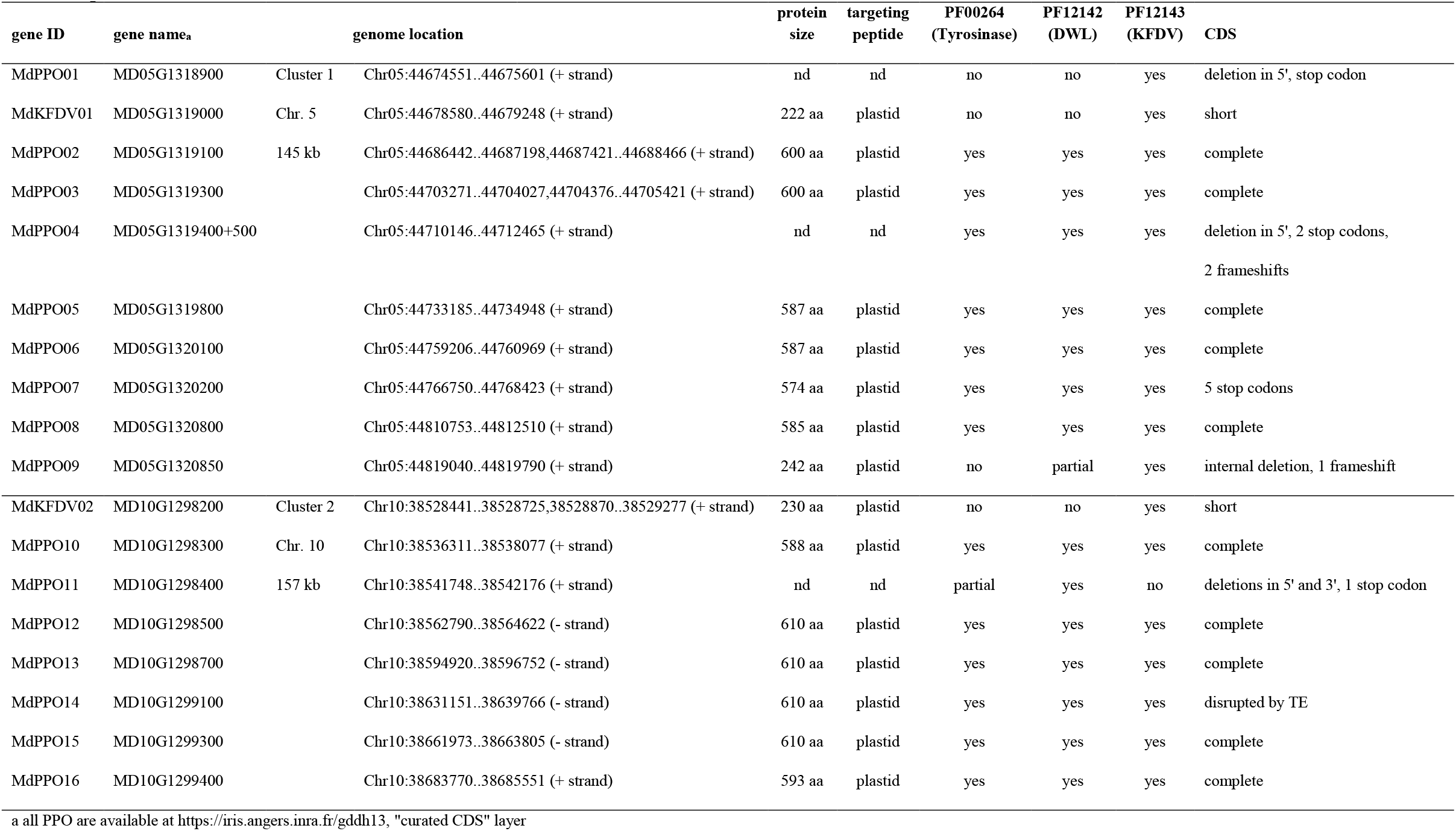
Ten *PPO* genes and two *PPO*-like genes in *Malus x domestica* ‘Golden Delicious’ double haploid 13. Chr: chromosome; nd: not determined; aa: amino acid; TE: transposable element

### Apple *PPO* gene expression profiles

To identify *PPO* promoters differentially responding to *Ea* infection, we quantified by RT-qPCR the specific expression of *PPO* genes in apple infected tissues. For this study pseudogenes (*MdPPO1*, *MdPPO4*, *MdPPO7*, *MdPPO9*, *MdPPO11* and *MdPPO14*) were discarded, as well as *MdPPO12*, *MdPPO13* and *MdPPO15* for which the design of specific primers was attempted repeatedly base on SNPs (Single Nucleotide Polymorphism; high level of identity ≥ 97.7 %; Online Resource 4), but failed. Primers for the remaining seven *PPO* and the two *PPO*-like genes were designed with the aim of quantifying their expression in *Ea* infected leaves of two apple genotypes with contrasted susceptibilities to fire blight, the susceptible ‘MM106’ and the resistant ‘Evereste’ (Venisse et al., 2002). During the test of primers efficacy performed using as template a cDNA pool from these *Ea* infected apple genotypes, we obtained very weak amplifications for *MdPPO02*, *MdPPO03*, *MdPPO05*, *MdPPO06*, *MdPPO08* and *MdPPO10* contrasting with the substantial ones for *MdKFDV01*, *MdKFDV02* and *MdPPO16* (Online Resource 5). Therefore gene expression kinetics are only shown for *MdKFDV01*, *MdKFDV02* and *MdPPO16* (Fig. 2). Analyses were performed in untreated leaves and in leaves challenged either with a wild-type strain of *Ea* (*Ea* wt) or a T3SS deficient mutant (*Ea* t3ss) or mock at 6, 24 and 48 hours post-treatment (hpt). A higher constitutive expression in untreated leaves of *MdPPO16* and *MdKFDV02* compared to *MdKFDV01* was observed in ‘Evereste’. *Ea* t3ss and mock treatments triggered similar expression changes in the two genotypes, peaking at 6 hpt especially for *MdPPO16* probably due to the stress caused by the infiltration method. A strong increase in *MdPPO16* expression was recorded in both genotypes challenged with *Ea* wt, suggesting a type III effector dependent induction. No noticeable modulation was observed in *MdKFDV01* and *MdKFDV02* expression levels whatever the treatment, except for *Ea* wt that seemed to slightly modulate the expression of *MdKFDV01* in ‘MM106’ at 24 and 48 hpt in one replicate only. Promoter of *MdKFDV02* from ‘MM106’, thereafter named *pKFDV02*, was selected for further investigation instead of promoter of *MdKFDV02* from ‘Evereste’ because expression of *MdKFDV02* was more stable throughout the kinetics (Fig. 2). Promoter of *MdPPO16* from ‘MM106’, thereafter named *pPPO16*, was also selected for further investigation instead of promoter of *MdPPO16* from ‘Evereste’ because *MdPPO16* expression throughout the kinetics was similar for the two genotypes (Fig. 2). We found 95.17 % identity between sequences of 2218 bp length upstream *MdPPO16* CDS in ‘MM106’ and ‘Evereste’.

**Fig. 2.**
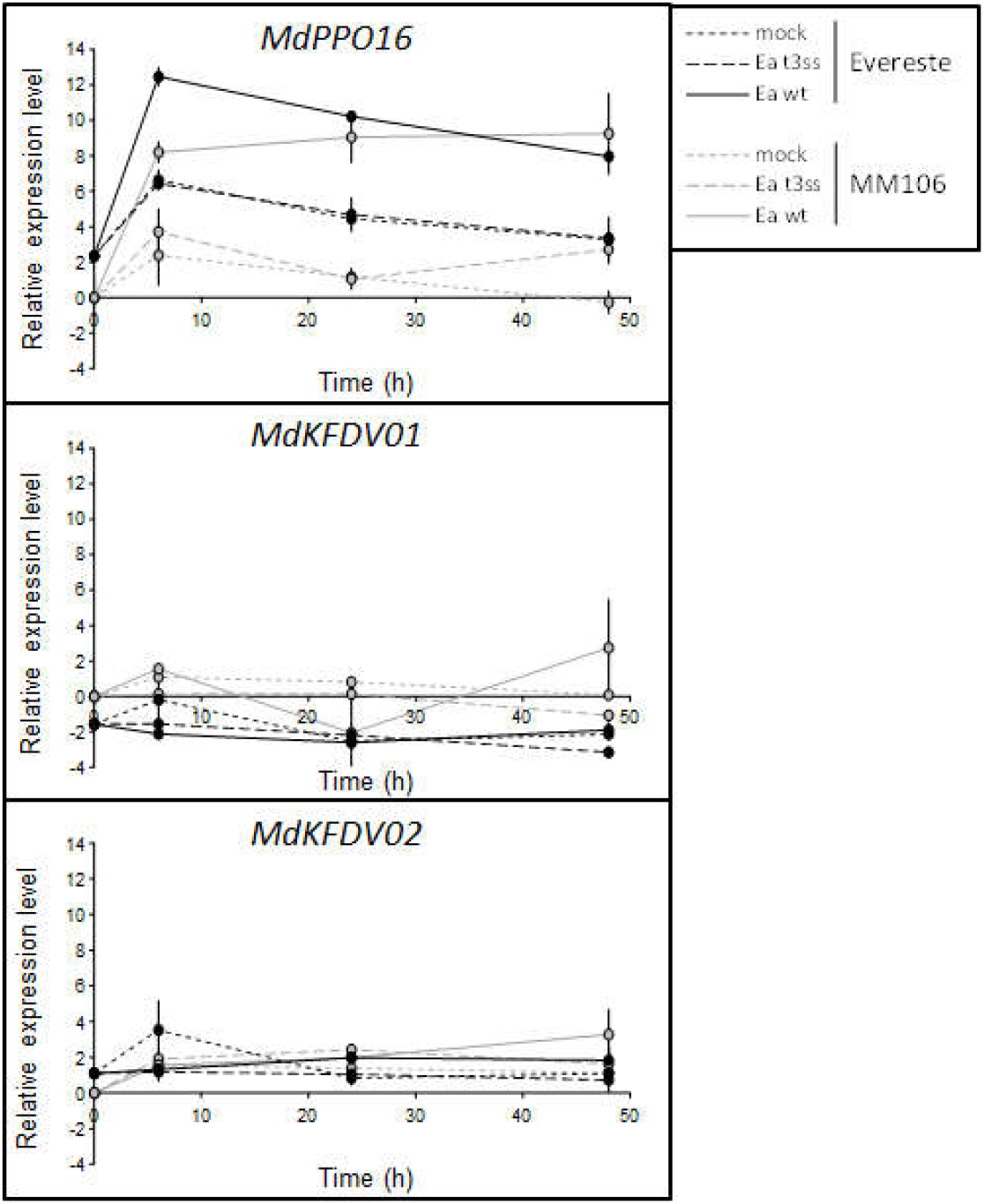
Expression profiling *MdKFDV01*, *MdKFDV02* and *MdPPO16* in ‘MM106’ (susceptible to fire blight) and ‘Evereste’ (resistant to fire blight) genotypes. Log_2_ expression levels were measured in untreated leaves and in mock, *Ea* t3ss or *Ea* wt infiltrated-leaves at 6, 24, 48 hpt. Expression levels for each gene are expressed relatively to the corresponding mean expression level of the untreated MM106 samples and normalized with 3 reference genes (*GAPDH*, *TuA* and *ACTIN*). Bars represent maximum and minimum values from two independent experiments (n=2)

### Promoter activity during transient expression

The regions upstream of *MdPPO16* (1177 bp; MK873007 in GenBank repository) and *MdKFDV02* (2030 bp; MK873006 in GenBank repository) CDS in ‘MM106’ genotype were cloned, and tested as a first approach in a transient expression assay in apple leaves using *GUS* (β-glucuronidase) as a reporter to quantify promoter activity in untreated, mock or *Ea*-infiltrated tissues. Rooted *in vitro* plants of ‘Gala’ were agroinfiltrated with EHA105 carrying different T-DNAs including *pPPO16:GUS*, *pKFDV02:GUS* or *p35S:GUS* as a control. Five days later, plants were infiltrated with mock or *Ea* wt and gene expression of *GUS* measured 24 hours later by RT-qPCR and calibrated to eliminate expression differences due to bacteria. *GUS* gene expression was stable in all samples under the control of *pKFDV02* (Fig. 3) and had comparable levels to that observed in *Ea*-infiltrated leaves under the control of *pPPO16*. Under the control of *pPPO16*, a strong induction of the *GUS* expression was observed in leaves challenged with *Ea* wt (a 5-fold increase approximately, Fig. 3). The same transient expression assay was repeated once in the other genotype ‘Golden Delicious’ with firefly luciferase (FIRE) instead of *GUS* as a reporter gene (Online Resource 6), to quantify promoter activity both at the transcriptional and enzymatic level. FIRE gene expression and protein activity were stable in all samples under the control of *pKFDV02* (Online Resource 6A and 6B respectively) and had comparable levels to that observed in untreated and mock-infiltrated leaves under the control of *pPPO16* or *p35S*. Under the control of *pPPO16*, a strong induction of the FIRE activity was observed in leaves challenged with *Ea* wt, both at the transcriptional and enzymatic level (a 2-fold increase approximately, Online Resource 6A and 6B).

**Fig. 3.**
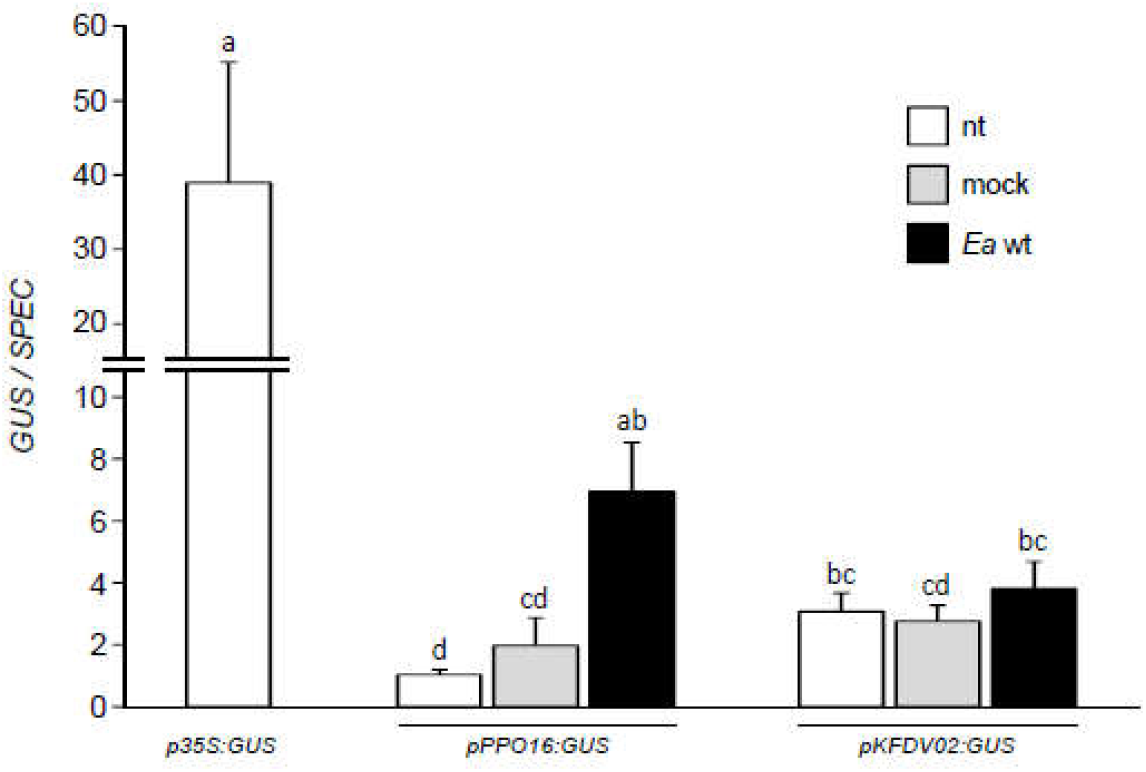
Expression of *GUS* gene driven by *pPPO16* and *pKFDV02* in transient assays. Relative expression of *GUS* reporter gene driven by *p35S*, *pPPO16* and *pKFDV02* in untreated (nt, white), mock (light gray) or *Ea* wt (black) -infiltrated leaves (24 hpt) of transiently transformed ‘Gala’ *in vitro* plants, six days after agroinfiltration. GUS raw expression levels of each sample were calibrated to the corresponding mean value of the sample *pPPO16:GUS-* nt, and normalized with *SPEC* gene to eliminate expression differences due to bacteria. Bars represent SEM from 3 biological repeats (n=3). Letters indicate statistical classes (Kruskal Wallis, p < 0.05)

### Promoter activity in stable transgenic clones

The contrasting results obtained with the transient assay encouraged us to perform apple stable transformations with two constructs carrying each promoter fused with the *GUS* gene as marker gene (*pPPO16:GUS* and *pKFDV02:GUS*), and to compare these to *p35S:GUS* transformed control. We respectively obtained one, two and four transgenic lines of ‘Golden Delicious’ transformed with *pPPO16:GUS, pKFDV02:GUS* and *p35S:GUS*. The unique line *pPPO16:GUS* and the more vigorous line *pKFDV02:GUS* were kept for subsequent analyses. For *p35S:GUS*, subsequent analyses were performed on two lines harboring a medium *GUS* expression (lines 217O and S; Online Resource 7). Assessment of transgenic lines selected for the further analyses are displayed in Online Resource 8. After *in vitro* multiplication, all stable transgenic lines were acclimatized and grown in greenhouse. The expression of the reporter gene was assessed by RT-qPCR in untreated, mock and *Ea* wt-infiltrated leaves at 24 hpt. In *pKFDV02* lines, activity was not significantly different from *p35S* lines in all conditions (nt, mock and *Ea* wt treatments, Fig. 4). By contrast, *GUS* expression was very weak in untreated and mock-infiltrated leaves of *pPPO16:GUS* lines and exhibited a strong and significant 10-fold induction in inoculated ones, reaching levels similar to *p35S:GUS* lines. Altogether these results corroborate those of the transient expression assay and show that *pPPO16* but not *pKFDV02* is strongly induced by *Ea* infection. The *pPPO16*-driven induction of the GUS in leaves challenged with *Ea* wt was also confirmed at the enzymatic level 40 and 48 hpt (Fig. 5).

**Fig. 4.**
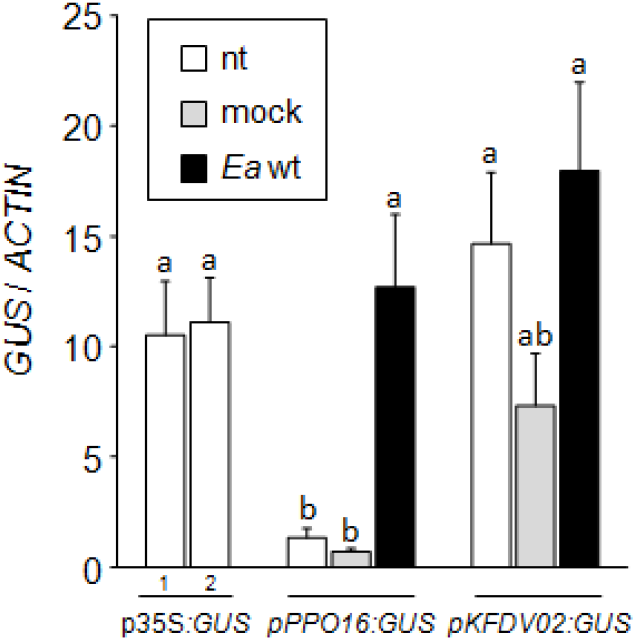
*pPPO16* and *pKFDV02*-driven *GUS* expression in ‘Golden Delicious’ transgenic lines cultivated in greenhouse and challenged with *Ea*. Relative expression of *GUS* reporter gene driven by *p35S, pPPO16* and *pKFDV02* in untreated leaves, mock or *Ea* wt-infiltrated leaves (24 hpt) from transgenic lines. *GUS* raw expression levels of each sample are relative to the corresponding mean value of the sample *pPPO16:GUS-nt*, and normalized with *ACTIN*. Numbers 1 and 2 represent independent lines of *p35S:GUS*. Bars represent SEM from 3 biological repeats (n=3). Letters indicate statistical classes (Kruskal Wallis, *p* < 0.05)

**Fig. 5.**
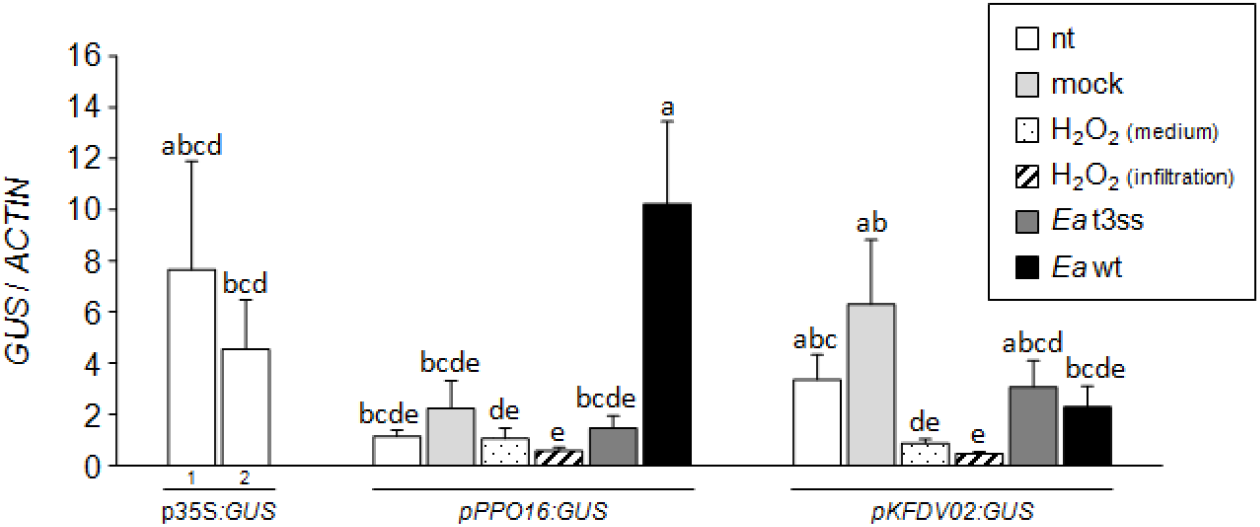
*pPPO16*-driven *GUS* activity in ‘Golden Delicious’ transgenic line challenged with *Ea*. Enzymatic activities of *GUS* reporter driven by *pPPO16* in untreated (nt, white), mock (light gray) and *Ea* wt (black) -infiltrated leaves of *in vitro* plants at 24, 40 and 48 hpt. *GUS* activity is expressed in nmoles MU (methylumbelliferone)/min/mg of total proteins. Bars represent SEM from 3 biological repeats (n=3). Letters indicate statistical classes (Kruskal Wallis, *p* < 0.05)

To determine which component of the *Ea* pathogenesis is responsible for the induction of *pPPO16*, i.e. a functional T3SS of the bacterium and/or the ROS production during the infectious process, *GUS* expression was recorded in transgenic rooted *in vitro* plants carrying *pPPO16:GUS* and *pKFDV02:GUS* at 24 hpt after the following different treatments: mock, *Ea* t3ss and *Ea* wt by leaf infiltration and H_2_O_2_ by leaf infiltration or by incorporation in the culture medium (Fig. 6). *GUS* expression was relatively stable when mediated by the promoter *pKFDV02*, although a slight but significant decrease of activity was observed after H_2_O_2_ treatments (infiltration and culture medium) compared to mock treatment. No change in *GUS* expression was observed in *pPPO16:GUS* line treated with mock, *Ea* t3ss and H_2_O_2_, while again a strong and significant 10-fold induction was observed when this line was inoculated with *Ea* wt. Taken together, these results highlight the ability of *Ea* to strongly and specifically induce *pPPO16* (and not *pKFDV02*), probably as an effect of a functional T3SS rather than H_2_O_2_ production.

**Fig. 6.**
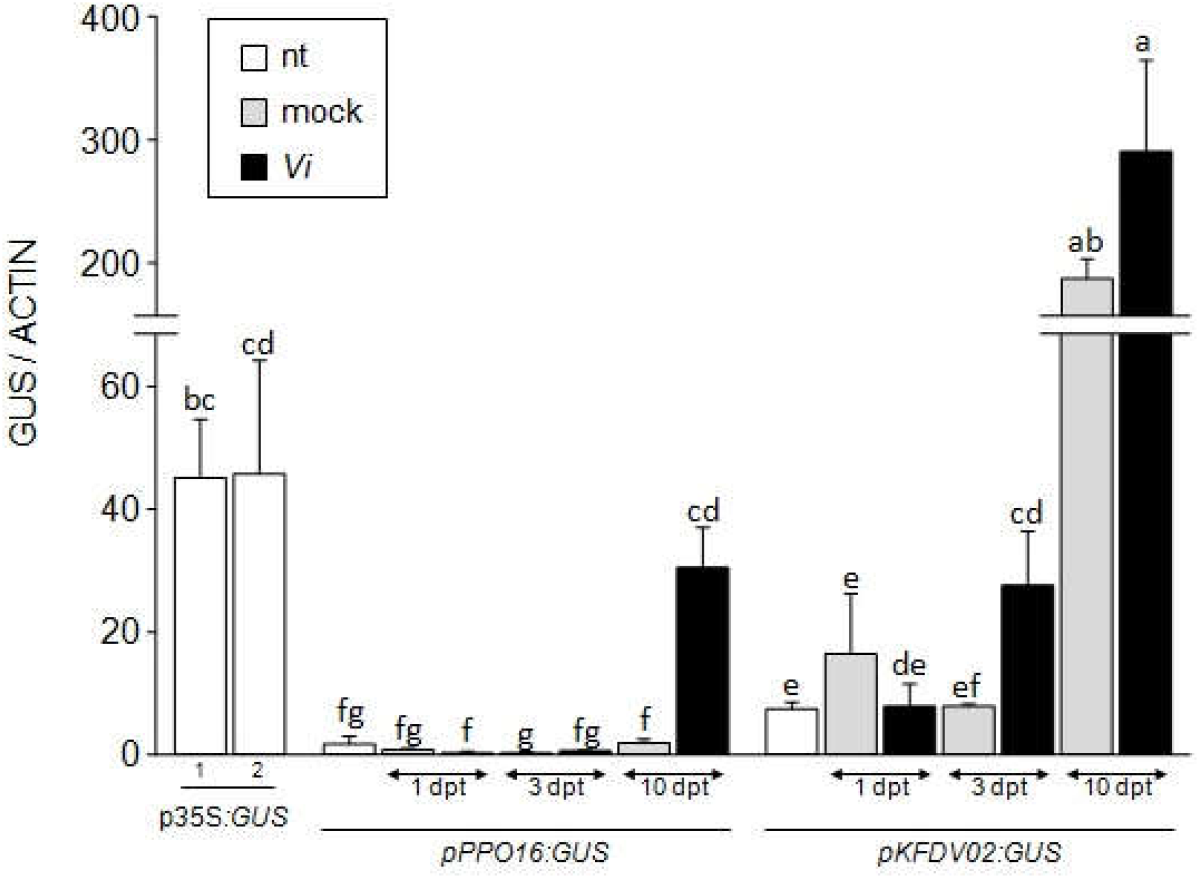
*pPPO16* and *pKFDV02-driven GUS* expression in ‘Golden Delicious’ transgenic lines challenged with *Ea*. Relative expression of *GUS* reporter driven by *p35S*, *pPPO16* and *pKFDV02* in untreated (nt), H_2_O_2_ -medium, mock or H_2_O_2_ or *Ea* t3ss or *Ea* wt-infiltrated leaves (24 hpt) from *in vitro* plants of transgenic lines. *GUS* raw expression levels of each sample are relative to the corresponding mean value of the sample *pPPO16:GUS*-nt, and normalized with *ACTIN*. Numbers 1 and 2 represent independent lines of *p35S:GUS*. Bars represent SEM from six biological repeats (n=6). Letters indicate statistical classes (Kruskal Wallis, *p* < 0.05)

In order to check *pPPO16* ability to be specifically activated by *Ea* and to observe *pKFDV02* behavior in response to another pathogen, the same transgenic lines were challenged with the pathogenic fungus *Vi* responsible for apple scab. Transgenic lines were therefore cultivated in greenhouse and *GUS* expression was assessed in untreated, mock and *Vi*-sprayed leaves at 1, 3 and 10 days post-treatment (dpt), the development of fungus being slower than that of *Ea*. Results indicated that up to 3 dpt, the *GUS* expression mediated by *pPPO16* was not affected by *Vi* in comparison to the corresponding mock controls (Fig. 7). However a strong and significant 15-fold induction was observed at 10 dpt, suggesting that *pPPO16* could be activated by another apple pathogen. Regarding *pKFDV02, GUS* expression was not significantly induced by *Vi* inoculation in the first 3 days, but considerably raised at 10 dpt in both mock or *Vi*-sprayed leaves, approximately 20-fold relative to the beginning of the experiment (*pKFDV02:GUS*-nt). The same phenomenon was also observed at 10 dpt in the youngest leaf of each plant which did not receive any treatment (Online Resource 9), suggesting the presence of a different unknown factor affecting *pKFDV02*.

## Discussion

Our work identified ten potentially functional apple PPO-encoding genes harboring the three known typical domains tyrosinase (PF00264), DWL (PF12142) and KFDV (PF12143), located on two duplicated chromosomes (5 and 10), all being addressed to the chloroplast and distributed in five phylogenetic sub-groups. This result completes the survey that Tran et al. (2012) performed among 25 land plants, describing *PPO* gene families varying in size (1 to 13) except in the genus *Arabidopsis* whose genome does not contain *PPO* sequences. Clustering of *PPO* genes at the same chromosomal location has already been observed in other plant species and indicates tandem gene duplications (Tran et al. 2012).

In the same chromosomal regions, we also identified six pseudogenes with similarity to *PPO* but with discrepancies such as deletions, premature stop codons and/or frameshifts, and two *PPO*-like genes of unknown function with only the KFDV domain. Doubts can be raised over their function as true polyphenol oxidases considering that they lack the common central domain of tyrosinase responsible of the oxidation process. Despite these doubts, KFDV genes were conserved in our study as *PPO-like* genes according to the fact that they have homologs in numerous dicot species.

Plant *PPO* genes are known to be involved in different physiological processes, from stress response to developmental regulation and environmental adaptation, as confirmed by their differential expression patterns in different situations (Thipyapong and Steffens, 1997; Constabel and Barbehenn, 2008; Tran and Constabel, 2011; Thipyapong et al. 2007). This makes regulatory sequences of *PPO* genes good candidates for diversified strategies of intragenesis. Unfortunately in our experiments, further analyses showed that the expression driven by *pKFDV02*, originally chosen for an expected constitutive activity was in fact modulated by unspecified factors. This result invalidated *pKFDV02* as a good candidate to drive a constitutive but weak expression for apple intragenesis development. On the other hand the fact that we found differential expression of *PPO* genes in response to *Ea* is coherent with previous works in other plant species showing induction in response to biotic stresses only for some *PPO* genes, in both incompatible and compatible interactions (Tran and Constabel, 2011; Rinaldi et al. 2007). In our hands *MdPPO16* induction in response to *Ea* has been recorded in three different genotypes (‘MM106’, ‘Evereste’ and ‘Gala’; Vergne et al. 2014 and this work). *MdPPO16* was also shown to be induced by wounding (Boss et al. 1995) and in fruit flesh browning disorder (Di Guardo et al. 2013), suggesting various functions for this gene.

Transient and stable transgenic assays using reporter genes fused to the immediate upstream region from the start codon of *MdPPO16* confirmed that this regulatory sequence was efficient to obtain the desired *Ea*-inducible expression pattern. Only one stable transgenic line was recovered with the *pPPO16-GUS* construction so we cannot affirm that the observed expression profile in that line is not affected, positively or negatively, by insertion effects. Despite this drawback, *pPPO16* promoter in 224C line show a quick and strong induction in leaves challenged with *Ea*, at the transcriptomic and enzymatic levels, in accordance with results get in transient assays with *GUS* or *FIRE* reporter genes. Thus we are confident on other results get with this line. In an intragenesis strategy designed to confer resistance to *Ea*, the use of such a promoter should ensure the precise induction of the intragene from the beginning of the infection process. Because a functional bacterial T3SS was required for this promoter induction, it should also avoid inappropriate activation in response to MAMPs (Microbial Associated Patterns, Choi and Klessig, 2016) of *Ea* or of other bacteria with similar conserved motifs potentially present on or inside the plant.

Induction of *pPPO16* seems to be linked to the loss of cellular integrity. Three lines of evidence support this hypothesis: (i) *pPPO16* induction requires *Ea* with a functional T3SS, which enables the injection of the major effector DspA/E into the plant cell, causing cell death (Boureau et al. 2006), (ii) *pPPO16* activation in compatible interaction with *Vi* occurred at 10 dpt in our experiments, which correspond to the beginning of tissue rupture by conidiogenesis (Ortega et al. 1998), and (iii) previous work shows the induction of *MdPPO16* after wounding (Boss et al. 1995). A specific induction of *pPPO16* linked to cell death is particularly interesting in the objective of controlling fire blight disease. It should ensure the induction of the intragene not only in the case of a real bacterial attack but also as a preventive barrier at wound sites caused by insects or climatic events, both acting as entry points for the bacteria. Despite the strong induction of *pPPO16* in response to *Vi* infection, it seems however unwise to consider this promoter in intragenic strategies for apple scab control, as it is only activated during the late phase of infection, i.e. conidiogenesis. Induction of a *PPO* gene during urediospore formation was already noticed in hybrid poplar / *Melampsora laricipopulina* interaction (Tran and Constabel, 2011).

We did not observe any response of *pPPO16* following exogenous application of H_2_O_2_, known as a precocious ROS produced during the oxidative burst accompanying Ea infection process (Vrancken et al. 2013). The concentration of H_2_O_2_ used in that work is moderate and known to modulate several defense genes in apple without leading to impaired tissue integrity (Dugé de Bernonville et al. 2014). The non-response of *pPPO16* following that moderate treatment should indicate that the expression driven by this promoter will remain stable despite moderate increase of H_2_O_2_ concentrations known to occur in various stress conditions (Saxena et al. 2016).

In the search for apple resistance, several cisgenic strategies have already been developed (Kost et al, 2015; Krens et al. 2015), but only one case of intragenic strategy has been tested, against another pathogen than *Ea* (*Vi*; Joshi et al. 2011). In order to create efficient fire blight intragenic resistances in apple, several candidate genes could be placed under the control of the *pPPO16* promoter characterized in our study: important regulators of defense pathways like *NPR1* (Malnoy et al. 2007), members of calcium-dependant protein kinases family (Kanchiswamy et al. 2013), genes involved in the jasmonic acid pathway (Dugé de Bernonville et al. 2012) or genes that increased oxidation of phenolic compounds (Flachowsky et al. 2010; Gaucher et al. 2013; Hutabarat et al. 2016).

The present work represents the first step towards the development of efficient “all native” solutions for apple fire blight resistance. As far as we know, *pPPO16* is the first cloned apple promoter inducible by *Ea*. Considering the narrowness of the gene pool screened to retrieve it, i. e. the *MdPPO* family, *pPPO16* could not be the best *Ea* inducible promoter candidate and comprehensive genomic level expression analyses are needed to find such candidates. Further work will be also needed to choose optimal candidate genes combining high efficiency for disease resistance, limited risk of break-down and absence of adverse effects on plant physiology.

## Supporting information

online resources 1 to 9

## Acknowledgements

The plasmid pGREEN II 0800-LUC was kindly provided by Dr. A. Allan (PFR, New Zealand). The plasmid pKGWFS7-35S was kindly provided by J. Jeauffre (IRHS, France). The authors gratefully acknowledge the IRHS-ImHorPhen team and the experimental unit HORTI of INRAE Angers for technical assistance in plant maintenance, B. Billy (SNES-GEVES) for technical assistance in flow cytometry and the technical platform ANAN. Technical contributions from M. Jacq, J.G. Bertault and P. Berthelot are also gratefully acknowledged. The authors wish to thank their collaborator Alexandre Degrave for his careful and critical reading of the manuscript.

## Statements and Declarations

### Funding

This project was funded by the INTRAPOM Project (INRAE BAP division) and post-doctoral grants from Region Pays de la Loire and Angers Agglomération (E. Vergne, L. Righetti and M. Gaucher).

### Competing interests

The authors have no relevant financial or non-financial interests to disclose.

### Author’s contribution

M.G. and L.R. were the main investigators in this study. They performed most of the experiments, analyzed and interpreted data, drafted the manuscript and revised it. E.V. designed the study, performed part of the experiments, analyzed and interpreted data, drafted the manuscript and revised it. S.A., T.D.B., M.N.B. and E.C. actively contributed to the analysis and interpretation of data and revised the manuscript. All authors read and approved the final version.

### Data availability

*pKFDV02* (MK873006) and *pPPO16* (MK873007) sequences are available in GenBank repository. Accession numbers of other sequences analyzed in this work;from repository GenBank or https://iris.angers.inra.fr/gddh13 (choose “Visit the apple genome”, enter accession number to search, right-click of mouse on the CDS of the “curated CDS” layer and choose view details to access sequence); are in Table 1, Online Resource 2 and 3, or in references given in these tables. Data generated during and/or analysed during the current study are available from the corresponding author on reasonable request.

## Figure legends

Fig. 7 *pPPO16* and *pKFDV02-driven GUS* expression in ‘Golden Delicious’ transgenic lines cultivated in greenhouse and challenged with *Vi*. Relative expression levels of *GUS* reporter gene driven by *p35S*, *pPPO16* and *pKFDV02* in untreated (nt), mock or *Vi*-sprayed leaves (1, 3, 10 dpt) from transgenic lines. *GUS* raw expression levels of each sample are relative to the corresponding mean value of the sample *pPPO16:GUS-nt*, and normalized with *ACTIN*. Numbers 1 and 2 represent independent lines of *p35S:GUS*. Bars represent SEM from 3 biological repeats (n=3). Letters indicate statistical classes (Kruskal Wallis, *p* < 0.05)

## Online Resources legends

**Online Resource 1** Pictures of some materials and methods. a) & b) *“In-vitro”* growing shoots 4 weeks after rooting. c) Vacuum chamber and pump used to infiltrate bacteria in leaves. d) Growing shoots of young grafts submerged in bacterial suspension to infiltrate bacteria in tissues by vacuum

**Online Resource 2** Agrobacterium strains used in this work

**Online Resource 3** Primers used in this work

**Online Resource 4** Percent identity matrix of CDS and protein sequences of PPO in *Malus* x *domestica*

**Online Resource 5** Mean Ct values obtained by RT-qPCR with specific primers designed on each coding sequence (CDS) and tested using a 4-fold serial dilution (from 1/16 to 1/4096) of a cDNA pool (all samples of ‘Evereste’ and ‘MM106’). Data were used to calculate primers efficiency and choose the genes for which the expression profiles were analyzed in the different samples (Fig. 2)

**Online Resource 6** Gene expression and activity of luciferase driven by *pPPO16* and *pKFDV02* in transient assays. Relative expression (A) and enzymatic activities (B) of firefly (FIRE) reporter driven by *p35S*, *pPPO16* and *pKFDV02* in untreated (nt, white), mock (light gray) or *Ea* wt (black) -infiltrated leaves (24 hpt) of transiently transformed ‘Golden Delicious’ *in vitro* plants, five days after agroinfiltration. *FIRE* raw expression levels (log2) of each sample were calibrated to the corresponding value of the sample *pPPO16:FIRE-nt*. Firefly luciferase expression and activity were normalized to Renilla luciferase (REN) expression and activity, respectively (n=1)

**Online Resource 7** *p35S*-driven *GUS* expression in four ‘Golden Delicious’ transgenic lines cultivated *in vitro*. Relative expression of *GUS* reporter gene driven by *p35S* in untreated leaves (nt) from transgenic lines 217F, O, R and S. *GUS* raw expression level of each sample are relative to the corresponding mean value in untreated leaves of the line 224C expressing *pPPO16:GUS*, and normalized with *ACTIN*. Bars represent SEM from 3 biological repeats (n=3). Lines 217O and S were kept for subsequent analyses

**Online Resource 8** Transgenic lines got are free from *A. tumefaciens* contamination. 217O & S: transgenic lines transformed with *p35S:GUS* construction, 222D: transgenic line transformed with *pKFDV02:GUS* construction, 224C: transgenic lines transformed with *pPPO16:GUS* construction, T+35S: DNA extraction of *A. tumefaciens* strain carrying pKGWFS7-*p35S:GUS* plasmid, as a positive control for transgenic lines transformed with *p35S:GUS* construction and *A. tumefaciens* presence, T+pKFDV02: DNA extraction of *E. Coli* strain carrying pKGWFS7-*pKFDV02:GUS* plasmid as a positive control for transgenic line transformed with *pKFDV02:GUS* construction, T+pPPO16: DNA extraction of *E. Coli* strain carrying pKGWFS7-*pPPO16:GUS* plasmid as a positive control for transgenic line transformed with *pPPO16:GUS* construction, NT: non-transformed ‘Gala’, T-:H20. *EF-1α, NptII*, *AGRO*, *p35S:GUS*, *pKFDV02:GUS*, *pPPO16:GUS*: primer couples

**Online Resource 9** Evolution of *GUS* expression over time in the youngest leaf of transgenic lines cultivated in greenhouse and challenged with *Vi*. Relative expression levels of *GUS* reporter gene promoted by *p35S*, *pPPO16* and *pKFDV02* in untreated leaves from seedlings of transgenic lines carrying the respective promoter in ‘Golden Delicious’ background. *GUS* raw expression levels for each sample are relative to the corresponding mean value of the sample *pPPO16:GUS-nt* (T0), and normalized with *ACTIN*. Numbers (1) and (2) represent independent lines of *p35S:GUS*. Bars represent SEM from 3 biological repeats (n=3). Letters indicate statistical classes (Kruskal Wallis, *p* < 0.05)

## Notes

### Competing Interest Statement

The authors have declared no competing interest.

### Summary of Updates

Results in Figure 3 have been revised to eliminate potential agrobacterium expression and new results have been added in a new Figure (actual Figure 5) to demonstrate GUS activity by fluorimetry(and not only expression) driven by pPPO16 promoter. Text has been improved.

