## Supplementary material for "An *Erwinia amylovora* inducible promoter for improvement of apple fire blight resistance": online resources 1 to 9

Online resource 1 Pictures of some materials and methods. a) & b) “*In-vitro*” growing shoots 4 weeks after rooting. c) Vacuum chamber and pump used to infiltrate bacteria in leaves. d) Growing shoots of young grafts submerged in bacterial suspension to infiltrate bacteria in tissues by vacuum

Online resource 2 Agrobacterium strains used in this work

| Strain |  | Vector |  | Resistance |  | Insert |  | Helper |  | Resistance |  | Use (concentration) |
| --- | --- | --- | --- | --- | --- | --- | --- | --- | --- | --- | --- | --- |
| EHA105 |  | pKGWFS7 |  | Spectinomycin 50 µg/mL |  | *p35S* |  | pBBR-MCS5 |  | Gentamicin  50 µg/mL |  | Transient or stable transformation |
|  |  | (Karimi et al., 2002) |  |  |  | *pKFDV02* (MM106; 2030 bp) |  |  |  |  |  | (1 x 10^9^ CFU/mL) |
|  |  |  |  |  |  | *pPPO16* (MM106; 2219 bp) |  |  |  |  |  |  |
|  |  | pBin61 |  | Kanamycin |  | *p35S:p19* |  | pBBR-MCS5 |  | Gentamicin  50 µg/mL |  | Transient transformation |
|  |  | (Bendahmane et al., 2000) |  | 50 µg/mL |  | (Voinnet et al., 2003) |  |  |  |  |  | (2.5 x 10^8^ CFU/mL) |
|  |  | pGREEN II 0800 LUC (Hellens et al., 2005) |  | Kanamycin |  | *p35S* (1027 bp) |  | pSoup |  | Tetracycline  10 µg/mL |  | Transient transformation |
|  |  |  |  | 50 µg/mL |  | *pKFDV02* (MM106 2030 bp) |  |  |  |  |  | (5 x 10^8^ CFU/mL) |
|  |  |  |  |  |  | *pPPO16* (MM106; 1177 bp) |  |  |  |  |  |  |

Online resource 3 Primers used in this work

| **Gene/family name** | **Accessions/ references** | **Forward primer 5'-3'** | **Reverse primer 5'-3'** | **Analyses** |
| --- | --- | --- | --- | --- |
| ***PPO and PPO-like genes*** | | | | |
| *MdPPO16* | MD10G1299400a | TGCCCGCCGCCTTCCAC | GCTCCATCGCTTTGTAGTATTTGTC | RT-qPCR |
| *MdKFDV01* | MD05G1319000a | TCCAAACAGGGTCGTCGAACT | TGCGCTTCCTTTTCCGCC |  |
| *MdKFDV02* | MD10G1298200a | CGTGCATGTGAACGATGATGAG | AACTCCCAAGTCCTCCAACAA |  |
| ***Promoters*** | | | | |
| *p35S* | from pK7WG2D from Karimi et al. (2002) | CACCACTAGAGCCAAGCTGATCTC | TCGACTAGAATAGTAAATTGTAAT | Cloning in pKGWFS7 via a pENTR/SD/D-TOPO |
|  |  | CACCGGTACCACTAGAGCCAAGCTGATCTC (KpnI) | CCCAAGCTTTCGACTAGAATAGTAAATTGTAAT (HindIII) | cloning in pGREEN II 0800 LUC |
| *pPPO16* | MK873007 | CACCGTTTCTTCATCCTGCTGCTT | GGCTTAGCTCTCTGGTTTTG | Cloning in pKGWFS7 via a pENTR/SD/D-TOPO |
|  |  | AAGGTACCTATCTGGCCAATTGCCTTGT (KpnI) | AACCATGGCTTAGCTCTCTGGTTTTG (NcoI) | cloning in pGREEN II 0800 LUC |
| *pKFDV02* | MK873006 | CACCGGTACCGAAGCTCAAGAAAACTGG (KpnI) | GGTTTTTGTTTCCTTTTTGTCAAG | Cloning in pKGWFS7 via a pENTR/SD/D-TOPO |
|  |  | CACCGGTACCGAAGCTCAAGAAAACTGG (KpnI) | AACCATGGTTTTTGTTTCCTTTTTGTCAAG (NcoI) | cloning in pGREEN II 0800 LUC |
| ***Reporter genes*** | | | | |
| *REN* (Renilla) | EU048863.1 Cloning vector pGreenII 0800 LUC | ATCGGACCCAGGATTCTTTT | ACTCGCTCAACGAACGATTT | RT-qPCR |
| *FIRE* (Firefly) | EU048863.1 Cloning vector pGreenII 0800 LUC | CCAGGGATTTCAGTC | AATCTCACGCAGGCAGTTCT |  |
| *GUS* | from pKGWFS7 from Karimi et al. (2002) | GCACGGGAATATTTCGCCAC | ATAACGGTTCAGGCACAGCA | specific RT (reverse) and RT-qPCR |
| *SPEC* | from pKGWFS7 from Karimi et al. (2002) | ATCATTCCGTGGCGTTATCC | GCTGGACCTACCAAGGCAAC |  |
| ***Reference genes*** | | | | |
| *GADPH* (Glyceraldehyde-3-phosphate dehydrogenase) | CN494000 | GCTGCCAAGGCTGTTGGAA | CAGTCAGGTCAACAACGGAAAC | RT-qPCR |
| *TuA* (Tubulin alpha-1) | CO065788 | GTTCAATGCTGTTGGTGGTG | CTGCGGAGAAGGATAGATGG |  |
| *ACTIN*(Actin 7) | CV151413 | CAACCTCTCGTCTGTGATAATG | GCATCCTTCTGTCCCATCC |  |
| *EF-1α* (Elongation Factor) | AJ223969 | CCTTCTTGAGGCTCTTGACCAG | CCAACAGGAACAGTACCGATACC | PCR |
| ***Transgenic lines assesment*** | | | | |
| *AGRO* | 23S ribosomal RNA coding gene CP014260.1 gene locus_tag="AWN88_17620" 1310643..1313449 | GTAAGAAGCGAACGCAGGGAACT | GACAATGACTGTTCTACGCGTAA | PCR |
| *NptII* | from plasmid pKGWFS7 from Karimi et al. (2002) | ATCGGGAGCGGCGATACCGTA | GAGGCTATTCGGCTATGACTG |  |
| *p35S:GUS* | promoter from pK7WG2D and reporter gene from pKGWFS7 from Karimi et al. (2002) | CGCACAATCCCACTATCCTT | ACAGTTTTCGCGATCCAGAC |  |
| *pKFDV02:GUS* | reporter gene from pKGWFS7 from Karimi et al. (2002) | AACCAATTGGGCTCGTGTAG | ACAGTTTTCGCGATCCAGAC |  |
| *pPPO16:GUS* | reporter gene from pKGWFS7 from Karimi et al. (2002) | CTGCGGGGTATCTACATGGT | TAATGAGTGACCGCATCGAA |  |
| a all PPO are available at https://iris.angers.inra.fr/gddh13, "curated CDS" layer | |  |  |  |

Online resource 4 Percent identity matrix of CDS and protein sequences of PPO in *Malus* x *domestica*

| CDS |  | **MdPPO02** | **MdPPO03** | **MdPPO05** | **MdPPO06** | **MdPPO08** | **MdPPO10** | **MdPPO12** | **MdPPO13** | **MdPPO15** | **MdPPO16** |
| --- | --- | --- | --- | --- | --- | --- | --- | --- | --- | --- | --- |
|  |  | **MD05G1319100** | **MD05G1319300** | **MD05G1319800** | **MD05G1320100** | **MD05G1320800** | **MD10G1298300** | **MD10G1298500** | **MD10G1298700** | **MD10G1299300** | **MD10G1299400** |
| **MdPPO02** | **MD05G1319100** | 100 | 91,5 | 66,65 | 66,07 | 66,07 | 61,71 | 61,98 | 61,8 | 62,37 | 63,74 |
| **MdPPO03** | **MD05G1319300** | 91,5 | 100 | 66,99 | 66,13 | 66,3 | 61,77 | 61,8 | 61,63 | 62,31 | 64,77 |
| **MdPPO05** | **MD05G1319800** | 66,65 | 66,99 | 100 | 94,1 | 93,67 | 63,83 | 62,52 | 62,11 | 61,88 | 64,15 |
| **MdPPO06** | **MD05G1320100** | 66,07 | 66,13 | 94,1 | 100 | 91,45 | 64,13 | 63,22 | 62,52 | 62,52 | 64,8 |
| **MdPPO08** | **MD05G1320800** | 66,07 | 66,3 | 93,67 | 91,45 | 100 | 63,52 | 61,4 | 61,23 | 60,94 | 64,61 |
| **MdPPO10** | **MD10G1298300** | 61,71 | 61,77 | 63,83 | 64,13 | 63,52 | 100 | 59,87 | 60,05 | 59,93 | 61,22 |
| **MdPPO12** | **MD10G1298500** | 61,98 | 61,8 | 62,52 | 63,22 | 61,4 | 59,87 | 100 | 98,31 | 97,71 | 74,29 |
| **MdPPO13** | **MD10G1298700** | 61,8 | 61,63 | 62,11 | 62,52 | 61,23 | 60,05 | 98,31 | 100 | 97,71 | 74,01 |
| **MdPPO15** | **MD10G1299300** | 62,37 | 62,31 | 61,88 | 62,52 | 60,94 | 59,93 | 97,71 | 97,71 | 100 | 73,89 |
| **MdPPO16** | **MD10G1299400** | 63,74 | 64,77 | 64,15 | 64,8 | 64,61 | 61,22 | 74,29 | 74,01 | 73,89 | 100 |
| Protein |  | **MdPPO02** | **MdPPO03** | **MdPPO05** | **MdPPO06** | **MdPPO08** | **MdPPO10** | **MdPPO12** | **MdPPO13** | **MdPPO15** | **MdPPO16** |
|  |  | **MD05G1319100** | **MD05G1319300** | **MD05G1319800** | **MD05G1320100** | **MD05G1320800** | **MD10G1298300** | **MD10G1298500** | **MD10G1298700** | **MD10G1299300** | **MD10G1299400** |
| **MdPPO02** | **MD05G1319100** | 100 | 92,49 | 61,14 | 60,62 | 60,66 | 51,58 | 54,25 | 54,08 | 54,93 | 56,55 |
| **MdPPO03** | **MD05G1319300** | 92,49 | 100 | 61,14 | 60,45 | 60,31 | 51,49 | 55,03 | 54,86 | 55,71 | 57,76 |
| **MdPPO05** | **MD05G1319800** | 61,14 | 61,14 | 100 | 93,36 | 91,79 | 54,77 | 54,43 | 53,74 | 52,52 | 57,17 |
| **MdPPO06** | **MD05G1320100** | 60,62 | 60,45 | 93,36 | 100 | 89,4 | 56,18 | 55,13 | 54,61 | 53,74 | 57,34 |
| **MdPPO08** | **MD05G1320800** | 60,66 | 60,31 | 91,79 | 89,4 | 100 | 56,03 | 54,01 | 53,66 | 52,61 | 56,39 |
| **MdPPO10** | **MD10G1298300** | 51,58 | 51,49 | 54,77 | 56,18 | 56,03 | 100 | 51,51 | 51,87 | 51,69 | 52,51 |
| **MdPPO12** | **MD10G1298500** | 54,25 | 55,03 | 54,43 | 55,13 | 54,01 | 51,51 | 100 | 97,54 | 96,23 | 70,97 |
| **MdPPO13** | **MD10G1298700** | 54,08 | 54,86 | 53,74 | 54,61 | 53,66 | 51,87 | 97,54 | 100 | 96,56 | 70,8 |
| **MdPPO15** | **MD10G1299300** | 54,93 | 55,71 | 52,52 | 53,74 | 52,61 | 51,69 | 96,23 | 96,56 | 100 | 70,63 |
| **MdPPO16** | **MD10G1299400** | 56,55 | 57,76 | 57,17 | 57,34 | 56,39 | 52,51 | 70,97 | 70,8 | 70,63 | 100 |

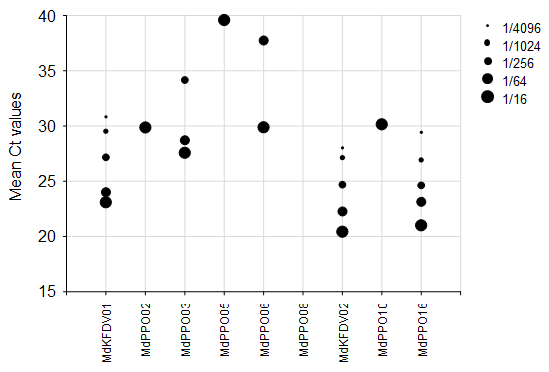

Online resource 5 Mean Ct values obtained by RT-qPCR with specific primers designed on each coding sequence (CDS) and tested using a 4-fold serial dilution (from 1/16 to 1/4096) of a cDNA pool (all samples of ‘Evereste’ and ‘MM106’). Data were used to calculate primers efficiency and choose the genes for which the expression profiles were analyzed in the different samples (Fig. 2)

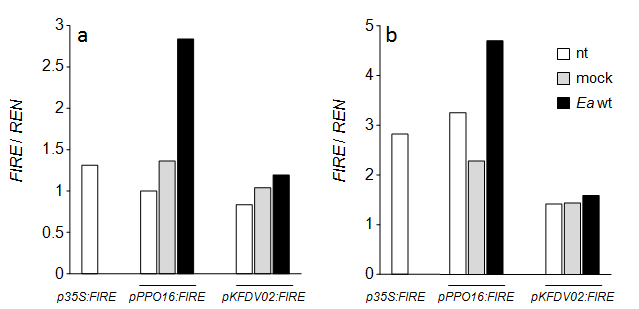

Online resource 6 Gene expression and activity of luciferase driven by *pPPO16* and *pKFDV02* in transient assays. Relative expression (A) and enzymatic activities (B) of firefly (FIRE) reporter driven by *p35S*, *pPPO16* and *pKFDV02* in untreated (nt, white), mock (light gray) or *Ea* wt (black) -infiltrated leaves (24 hpt) of transiently transformed ‘Golden Delicious’ *in vitro* plants, five days after agroinfiltration. *FIRE* raw expression levels (log2) of each sample were calibrated to the corresponding value of the sample *pPPO16:FIRE*-nt. Firefly luciferase expression and activity were normalized to Renilla luciferase (REN) expression and activity, respectively (n=1)

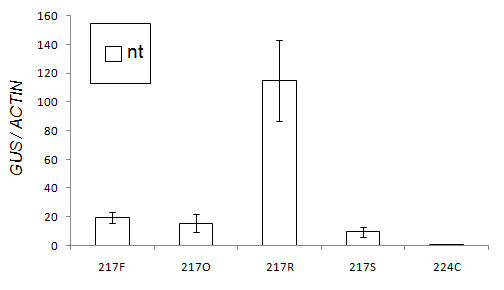

Online resource 7 *p35S*-driven *GUS* expression in four ‘Golden Delicious’ transgenic lines cultivated *in vitro*. Relative expression of *GUS* reporter gene driven by *p35S* in untreated leaves (nt) from transgenic lines 217F, O, R and S. *GUS* raw expression level of each sample are relative to the corresponding mean value in untreated leaves of the line 224C expressing *pPPO16:GUS*, and normalized with *ACTIN*. Bars represent SEM from 3 biological repeats (n=3). Lines 217O and S were kept for subsequent analyses

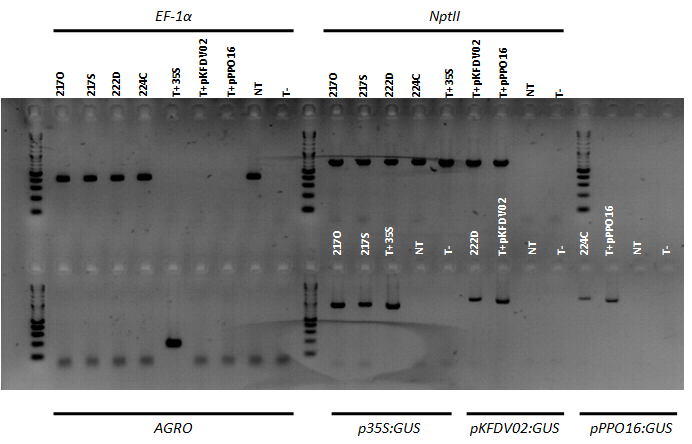

Online resource 8 Transgenic lines got are free from *A. tumefaciens* contamination. 217O & S: transgenic lines transformed with *p35S:GUS* construction, 222D: transgenic line transformed with *pKFDV02:GUS* construction, 224C: transgenic lines transformed with *pPPO16:GUS* construction, T+35S: DNA extraction of *A. tumefaciens* strain carrying pKGWFS7-*p35S:GUS* plasmid, as a positive control for transgenic lines transformed with *p35S:GUS* construction and *A. tumefaciens* presence, T+pKFDV02: DNA extraction of *E. Coli* strain carrying pKGWFS7-*pKFDV02:GUS* plasmid as a positive control for transgenic line transformed with *pKFDV02:GUS* construction , T+pPPO16: DNA extraction of *E. Coli* strain carrying pKGWFS7-*pPPO16:GUS* plasmid as a positive control for transgenic line transformed with *pPPO16:GUS* construction, NT: non-transformed ‘Gala’, T-:H20. *EF-1α*, *NptII*, *AGRO*, *p35S:GUS*, *pKFDV02:GUS*, *pPPO16:GUS*: primer couples

Online resource 9 Evolution of *GUS* expression over time in the youngest leaf of transgenic lines cultivated in greenhouse and challenged with *Vi*. Relative expression levels of *GUS* reporter gene promoted by *p35S*, *pPPO16* and *pKFDV02* in untreated leaves from seedlings of transgenic lines carrying the respective promoter in ‘Golden Delicious’ background. *GUS* raw expression levels for each sample are relative to the corresponding mean value of the sample *pPPO16:GUS*-nt (T0), and normalized with *ACTIN*. Numbers (1) and (2) represent independent lines of *p35S:GUS*. Bars represent SEM from 3 biological repeats (n=3). Letters indicate statistical classes (Kruskal Wallis, *p* < 0.05)
